# Distinct Patterns of SARS-CoV-2 BA.2.87.1 and JN.1 Variants in Immune Evasion, Antigenicity and Cell-Cell Fusion

**DOI:** 10.1101/2024.03.11.583978

**Authors:** Pei Li, Yajie Liu, Julia Faraone, Cheng Chih Hsu, Michelle Chamblee, Yi-Min Zheng, Claire Carlin, Joseph S. Bednash, Jeffrey C. Horowitz, Rama K. Mallampalli, Linda J. Saif, Eugene M. Oltz, Daniel Jones, Jianrong Li, Richard J. Gumina, Shan-Lu Liu

## Abstract

The rapid evolution of SARS-CoV-2 variants presents a constant challenge to the global vaccination effort. In this study, we conducted a comprehensive investigation into two newly emerged variants, BA.2.87.1 and JN.1, focusing on their neutralization resistance, infectivity, antigenicity, cell-cell fusion, and spike processing. Neutralizing antibody (nAb) titers were assessed in diverse cohorts, including individuals who received a bivalent mRNA vaccine booster, patients infected during the BA.2.86/JN.1-wave, and hamsters vaccinated with XBB.1.5-monovalent vaccine. We found that BA.2.87.1 shows much less nAb escape from WT-BA.4/5 bivalent mRNA vaccination and JN.1-wave breakthrough infection sera compared to JN.1 and XBB.1.5. Interestingly. BA.2.87.1 is more resistant to neutralization by XBB.15-monovalent-vaccinated hamster sera than BA.2.86/JN.1 and XBB.1.5, but efficiently neutralized by a class III monoclonal antibody S309, which largely fails to neutralize BA.2.86/JN.1. Importantly, BA.2.87.1 exhibits higher levels of infectivity, cell-cell fusion activity, and furin cleavage efficiency than BA.2.86/JN.1. Antigenically, we found that BA.2.87.1 is closer to the ancestral BA.2 compared to other recently emerged Omicron subvariants including BA.2.86/JN.1 and XBB.1.5. Altogether, these results highlight immune escape properties as well as biology of new variants and underscore the importance of continuous surveillance and informed decision-making in the development of effective vaccines.

## INTRODUCTION

Severe acute respiratory syndrome coronavirus 2 (SARS-CoV-2), the causative agent of the COVID-19 pandemic, continues to evolve despite the global pandemic being declared over. Late 2023 into early 2024 has seen the emergence of highly mutated variants of the virus, heightening new concern over the continued efficacy of current vaccination strategies and other pandemic control measures (1, 2). Among these, the BA.2.86 variant was characterized by around 30 mutations and evolved into JN.1 and a series of other subvariants with the spike protein distinct from the previously dominant variant XBB.1.5 (1). While BA.2.86 proved to be a less dominant variant and displayed minimal escape of neutralizing antibodies in mRNA-vaccinated and SARS-CoV-2 infected sera (3, 4), JN.1, which has only an additional L455S mutation in spike compared to BA.2.86, has significantly increased evasion of neutralizing antibodies and become the dominant variant in the United and States and other countries (5, 6).

Concern is mounting once more as a new highly mutated variant, BA.2.87.1, has been detected in South Africa (7). This variant contains over 100 mutations relative to XBB.1.5 and JN.1 throughout the genome, with over 30 in spike alone (**Fig. 1a**) (1). Since its initial detection in September of 2023, 9 cases have been recorded in South Africa as of early February 2024 and was recently reported in the wastewater of Southeast Asia. This variant has not yet been detected elsewhere (7). Currently, little is known about this new variant, including critical aspects of virus biology, sensitivity to neutralizing antibodies, and transmissibility. While BA.2.87.1 does not appear to have spread widely now, the fact that the currently dominant JN.1 was derived from a single mutation L455S in the spike in the less-fit BA.2.86 variant raises concerns over similar situations occurring.

**FIG 1.**
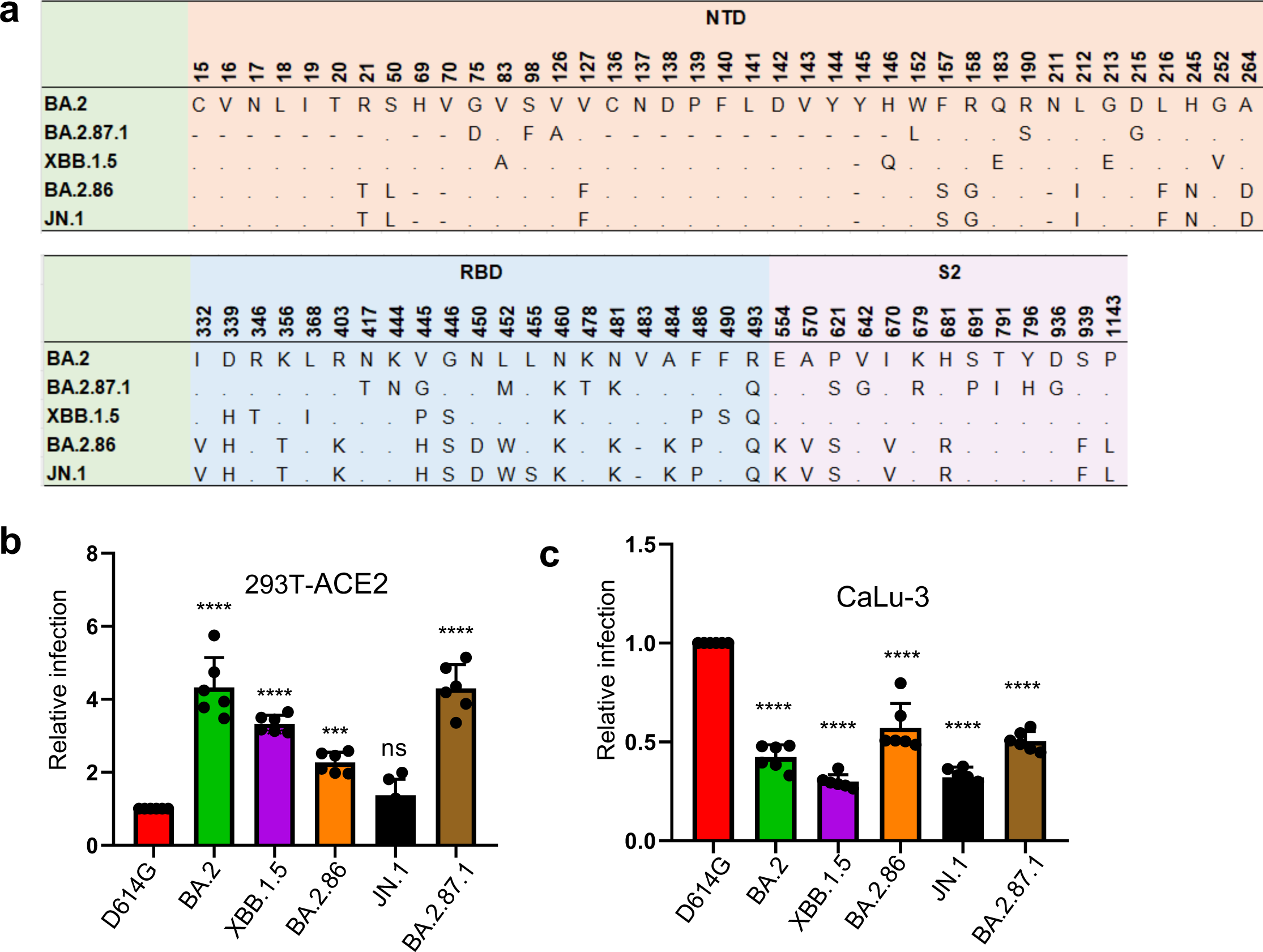
Infectivity of BA.2.87.1 and JN.1 in 293T-ACE2 and CaLu-3 cells. (a) A schematic depiction comparing spike mutations in the studied variants including BA.2.87.1 and JN.1 by amino acid numbers. NTD = N-terminal domain, RBD = receptor binding domain; S2: the S2 subunit region. (b-c) Relative infectivity of lentiviral pseudotypes bearing each of the listed spikes in (b) 293T cells expressing human ACE2 (293T-ACE2) and (c) human lung cell line CaLu-3. Relative luminescence readouts were normalized to D614G (D614G = 1.0). Bars in (b–c) represent means ± standard error from triplicates of transfection. Significance relative to D614G was analyzed by a one-way repeated measures ANOVA with Bonferroni’s multiple testing correction (n = 6). p values are displayed as ns p > 0.05, ***p < 0.001, and ****p < 0.0001.

Here, we investigate the immune escape and biology of the BA.2.87.1 variant in comparison to previously dominant variants JN.1 and XBB.1.5 and ancestral BA.2.86, BA.2 and parental D614G. We characterized the nAb titers in the sera of health care workers (HCWs) that received the wildtype (WT) plus BA.4/5 spike bivalent mRNA vaccine (n=13), sera from hamsters that received the XBB.1.5 monovalent mRNA vaccine (n=15), and sera from patients in the ICU during the BA.2.86/JN.1-wave of infection in Columbus, Ohio, U.S (n=9). We also elucidated the antigenic distance between variants and examined the neutralization of two RBD-targeting monoclonal antibodies S309 and 2B04. Additionally, we studied other aspects of virus biology including viral infectivity in lung airway epithelial cells, spike processing into the S1 and S2 subunits, spike surface expression, and cell-cell fusion.

## RESULTS

### BA.2.87.1 exhibits comparable infectivity to its ancestral BA.2 in human 293T-ACE2 and lung epithelial CaLu-3 cells

We first investigated the infectivity of pseudotyped lentiviral vectors bearing the spike of BA.2.87.1 or others of interest in 293T cells overexpressing human ACE2 (293T-ACE2) (**Fig. 1b**) and human lung epithelial cell line CaLu-3 (**Fig. 1c**). In 293T-ACE2 cells, BA.2.87.1 exhibited comparable infectivity to BA.2, but with a 4-fold increase relative to D614G (p < 0.0001). In contrast, JN.1 showed an infectivity comparable to D614G but lower than BA.2 (3.2-fold, p < 0.0001), BA.2.87.1 (3.1-fold, p < 0.0001) and XBB.1.5 (2.4-fold, p < 0.0001), respectively. The infectivity of JN.1 was even lower than its ancestral BA.2.86, with a 40% decrease (p < 0.01), and was among the lowest in all examined Omicron subvariants (**Fig. 1b**).

Omicron spikes have been characterized by an overall lower infectivity in CaLu-3 cells, but infectivity increased with some of the recently emerged subvariants (8–12). Here we found that both JN.1 and BA.2.87.1 had titers about 2-fold lower in relative infectivity compared to D614G (p < 0.0001), but 1.6-fold (p < 0.0001) and 1.7-fold (p < 0.0001) higher than JN.1 and XBB.1.5, respectively. Notably BA.2.86 showed an increased infectivity in CaLu-3 cells compared to other Omicron subvariants, similar to previous results (4, 13–15) (**Fig. 1c**).

### Bivalent mRNA-vaccinated sera more effectively neutralize BA.2.87.1 than JN.1

We next investigated the nAb responses in a series of cohorts (**Fig. 2, Fig. S1**). The first was The Ohio State University (OSU) Wexner Center HCWs that received at least 2 doses of monovalent vaccine (WT) plus a single booster of bivalent vaccine (WT + BA.4/5) (**Table S1**). The samples were collected between December 2022 and January 2023, approximately 23 and 108 days post the bivalent dose administration; the cohort had no breakthrough infection with BA.2.86/JN.1 or BA.2.87.1, but 9 of the 13 were COVID-19 positive with variants prior to the XBB.1.5 wave (see **Table S1**). BA.2.87.1 exhibited an increased sensitivity to neutralization by the bivalent mRNA-vaccine sera, with a titer ∼4-fold lower than D614G (p < 0.05) as compared to JN.1, which was 7.6-fold lower than D614G (p < 0.001) (**Fig. 2a, Fig. S1a**). JN.1 exhibited the lowest titers of all variants tested, even relative to its ancestral BA.2.86 and previous XBB.1.5, which were 4.7-fold and 4.8-fold lower than D614G (p < 0.05 for both), respectively. However, all variants were effectively neutralized by the bivalent HCW sera, with none falling below the limit of detection for the assay (NT_50_ = 40). These results together suggest that bivalent mRNA vaccine could still be effective for BA.2.87.1 but efficiency is reduced for JN.1.

**FIG 2.**
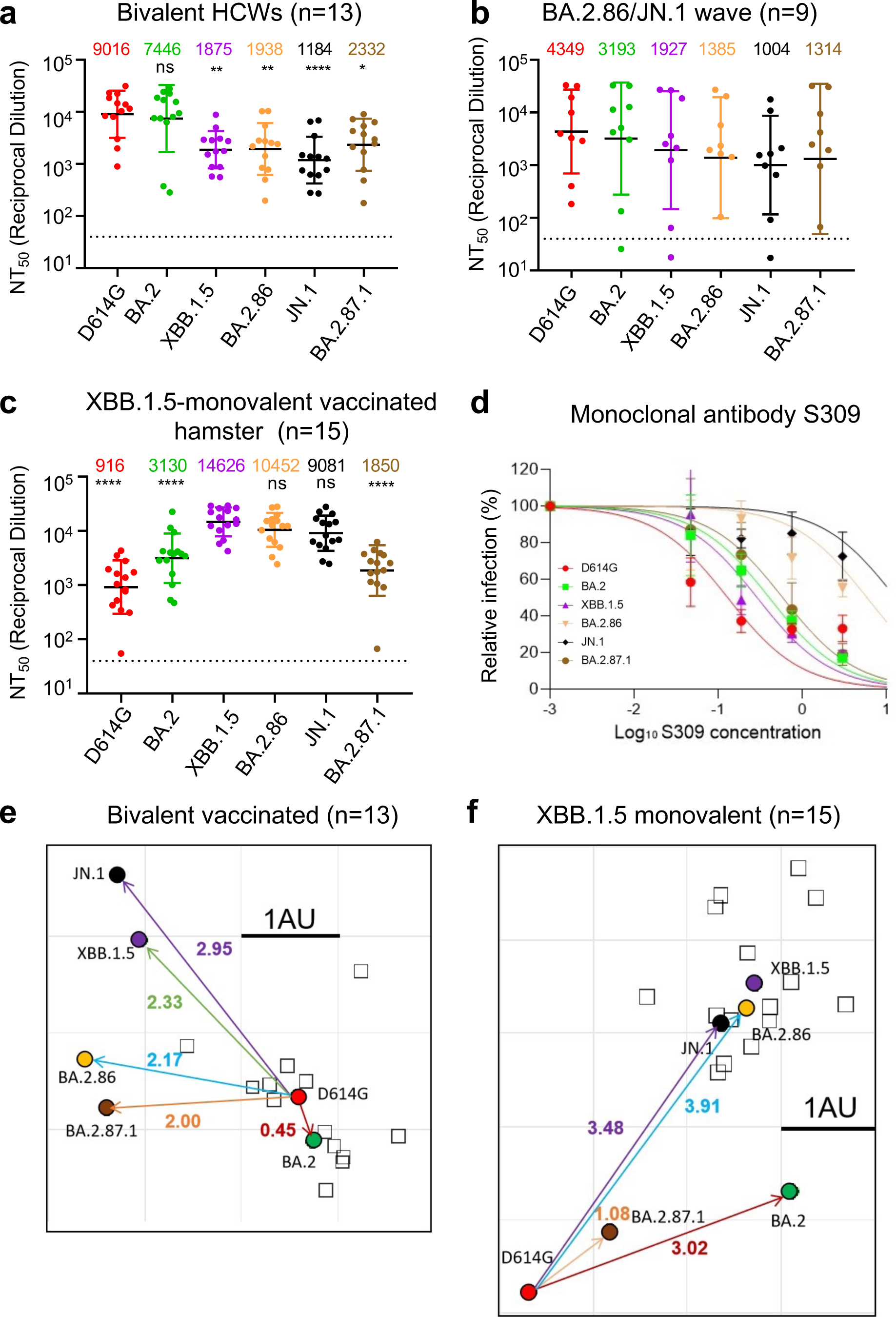
Neutralization of BA.2.87.1 and JN.1 by bivalent-vaccinated human sera, JN.1-wave human sera, XBB.1.5-vaccinated hamster sera, and monoclonal antibody S309. (a–c) NAb titers were determined using lentiviruses bearing the indicated spike proteins, with the titer of D614G as a control. All were compared against D614G or XBB.1.5 unless otherwise specified. The three cohorts included sera from 13 HCWs who had at least 2 monovalent doses of mRNA vaccine and 1 dose of bivalent mRNA vaccine (n = 13) (a), sera from Columbus first-responder/household contact cohort (P1 to P5) and ICU patients admitted to OSU Wexner Medical Center (P6 to P9) during when the BA.2.86/JN.1 variants were predominantly circulating in Columbus, Ohio (b) (n=9 total), and sera from Golden Syrian hamsters inoculated with two doses of XBB.1.5 monovalent vaccine (recombinant mumps virus expressing the spike of XBB.1.5, 1.5 x 10^5^ PFU per hamster, three weeks apart) (n=15), with blood being collected 5 weeks after inoculation (c). Geometric mean NT_50_ values for each variant are shown on the top. Bars represent geometric means with 95% confidence intervals. Statistical significance was analyzed with log10 transformed NT_50_ values. Comparisons between multiple groups were performed using a one-way ANOVA with Bonferroni post-test. Dashed lines represent the threshold of detection, i.e., NT_50_ = 40. p values are shown as ns p > 0.05, *p < 0.05, **p < 0.01, ****p < 0.0001. (d) Neutralization by mAb S309 was assessed, with representative plot curves displayed. Bars represent means ± standard deviation. (e–f) Antigenic maps for neutralization titers from Fig. 2a (bivalent-vaccinated human sera) and Fig. 2c (XBB.1.5-monovalent-vaccinated hamster sera) were made using the Racmacs program (1.1.35) (see Methods). Squares represent the individual sera sample and circles represent variants. One square on the grid represents one antigenic unit squared.

### Sera from JN.1/BA.2.86-wave ICU patients neutralize BA.2.87.1 better compared to JN.1 and XBB.1.5

The next cohort we investigated were Columbus first-responders and their household contacts (n=5, P1 to P5) as well as ICU COVID-19 patients admitted to the OSU Medical Center (n=4, P6 to P9) during the BA.2.86/JN.1 wave of infection in Columbus, OH (early 2024) (total n=9 in this cohort) (**Fig. 2b, Fig. S1b,** and **Table S1**). Nasal swabs were collected and sequenced, with 1 individual being confirmed to have been infected with BA.2.86, 1 individual confirmed to have been infected with JN.1, and the remaining 7 were assumed to have been infected with JN.1 based on the timing of the cases in Columbus, Ohio after Jan 2024. Of note, all nine patients were vaccinated with different doses of mRNA vaccine, most 357-898 days prior to sample collection, except one (P5), who was vaccinated with XBB.1.5 monovalent vaccine with sample collected 45 days after the vaccination (**Table S1**). Overall, nAb titers varied greatly in this cohort due to its heterogeneity, and were generally lower compared to the bivalent vaccinated cohort, especially against Omicron-lineage variants (**Figs. 2a-b** and **Figs. S1a-b**). Notably, BA.2.87.1 exhibited a modestly increased titer compared to JN.1 (1.3-fold, p = 0.301), with only 3.3-fold lower than D614G (p = 0.6778). Surprisingly, JN.1 showed the lowest neutralization titers, which were similar to the bivalent serum samples (**Fig. 2a**, **Fig. S1a**), with ∼4.3-fold lower than D614G (p = 0.1321). Notably, despite the limited sample size, 3 of the 4 ICU patients (P6, P8 and P9) exhibited very high neutralization titers compared to the first-responders and household contacts, results of which were in accordance with our previous studies (4, 9, 10, 13). We noticed that one ICU patient (P7, 78-year-old female) and one first-responder and household contact (P1) exhibited extremely low titers, especially against the Omicron variants (**Fig. 2b**, **Fig. S1b**). This was despite that P7 had received 4 doses of monovalent WT mRNA and 2 doses of WT-BA.4/5 bivalent vaccine shots prior to the BA.2.86/JN.1-wave in July 2023, without obvious history of immunocompromised conditions.

### BA.2.87.1 is less efficiently neutralized by XBB.1.5 monovalent-vaccinated hamster sera compared to JN.1

The final cohort we tested was a group of hamsters vaccinated twice with a monovalent XBB.1.5 spike vaccine delivered by recombinant mumps virus (n=15). In contrast to the human cohorts that received WT and BA.4/5 bivalent vaccine doses shown above, we found that these hamster serum samples exhibited the highest titers against XBB.1.5 (NT_50_ = 14,626), BA.2.86 (NT_50_ = 10,452), and JN.1 (NT_50_ = 9,081), with D614G showing the lowest titers (NT_50_ = 916), followed by BA.2.87.1 (NT_50_ = 1,850) and BA.2 (NT_50_ = 3,130) (**Fig. 2c, Fig. S1c**). For this cohort, comparisons were thus made instead to XBB.1.5 rather than D614G, due to the fact that XBB.1.5 is the variant included in the vaccine. Titers against JN.1 were only slightly reduced, with 1.6-fold lower than XBB.1.5 (p = 0.4722). Titers against BA.2.87.1 were markedly reduced, with 7.9-fold lower than XBB.1.5 (p < 0.0001). No neutralization escape was evident for this cohort relative to XBB.1.5, though one hamster (XBB.1.5-15) exhibited titers near the limit of detection for both D614G and BA.2.87.1 (**Fig. 2c, Fig. S1c**).

### Class III monoclonal antibody S309 efficiently neutralizes BA.2.87.1 but not JN.1

We next tested the neutralization of BA.2.87.1 and JN.1 by two neutralizing antibodies: the class III monoclonal antibody (mAb) S309 and class I mAb 2B04 (16, 17). S309 targets the epitopes of non-receptor binding motif (RBM) of the spike and has largely maintained efficacy against Omicron variants with the exception of CH.1.1, CA.3.1, BA.2.75.2 and BA.2.86 (9, 18). Interestingly, we found that S309 maintained neutralization against BA.2.87.1, with an IC_50_ of 0.62 µg/mL (**Fig. 2d, Fig. S1d**). However, the neutralizing activity of S309 was lost for JN.1 and greatly reduced for BA.2.86, with an IC_50_ of 6.22 µg/mL for the latter (**Fig. 2d, Fig. S1d**). Omicron variants have been expected to exhibit a complete escape of mAb 2B04 due to the multitude of mutations contained within the class I RBM epitope (1, 19) (**Fig. 1a**), and JN.1 and BA.2.87.1 were no exception, both having escaped neutralization by this monoclonal antibody (**Figs. S1d-e**).

### BA.2.87.1 is antigenically more related to D614G and BA.2 other than recent Omicron subvariants

To further analyze our neutralization data, we performed antigenic cartography analysis using a program called Racmacs, which uses principal component analysis to plot the antigenic distance between the variants tested based on the nAb titers. For bivalent-vaccinated human samples, D614G and BA.2 clustered near each other, with an antigenic distance of 0.45, and they were farther away from the cluster of newer variants (**Fig. 2e**). Notably, JN.1 was farthest away from D614G, with antigenic distance of 2.95, which was in accordance with its lowest nAb titers (**Fig. 2a** and **Fig. S1a**), suggesting that JN.1 is more antigenically distinct from D614G and BA.2 than XBB.1.5, BA.2.86, and BA.2.87.1. Interestingly, BA.2.87.1 clustered closer to D614G and BA.2, with an antigenic distance of 2 and 2.15., respectively, suggesting that despite the 30 additional mutations in the spike, it has actually become more antigenically similar to the parental variants (**Fig. 2e**). Because of the heterogeneity as well as the small sample size of JN.1-wave patient samples, we did not perform the antigenic analysis for this cohort.

The hamster cohort map was quite distinct from the bivalent mRNA-vaccinated human cohort due to the very different patterns of antigenic exposure. We observed that XBB.1.5, BA.2.86, and JN.1 all clustered together, but with greater antigenic distances of 3.48∼4.14 from D614G; whereas BA.2.87.1 was antigenically closer with distances of 1.08 and 2 from D614G and BA.2, respectively (**Fig. 2f**). Overall, these analyses indicate that antigenically, BA.2.87.1 is more closely related to BA.2, the ancestral Omicron variant; however, BA.2.86 and JN.1 are more closely related to XBB.1.5.

### BA.2.87.1 spike exhibits increased cell-cell fusion and processing into S1 and S2

Given more than 30 amino-acid changes in the spike protein of BA.2.87.1 and JN.1, including some near the furin cleavage site as well as in the S2 subunit (**Fig. 1a**), it is important to examine the furin cleavage efficiency and cell-cell fusion property of these new variants. For cell-cell fusion, we transfected 293T cells with the spikes of interest plus GFP, followed by co-culturing the detached effector 293T cells with target 293T-ACE2 or CaLu-3 cells. In both cell lines, D614G exhibited the highest cell-cell fusion compared to all Omicron variants (**Figs. 3a-d**), as would be expected. Notably, BA.2.87.1 exhibited the highest cell-cell fusion activity of the Omicron variants in both cell lines. While JN.1 exhibited an increased cell-cell fusion relative to BA.2, the level was comparable to its ancestral BA.2.86. XBB.1.5 showed increased fusion activity relative to the ancestral BA.2, which was consistent with our previous results (4, 9), although the level was relatively lower than BA.2.87.1 in both 293T-ACE2 and CaLu3 cells (**Figs. 3a-d)**. We validated these results using a syncytia formation assay wherein 293T-ACE2 cells are transfected to produce the spikes of interest and GFP and incubated 24 hours before imaging fusion (**Figs. S2a and b**).

**FIG 3.**
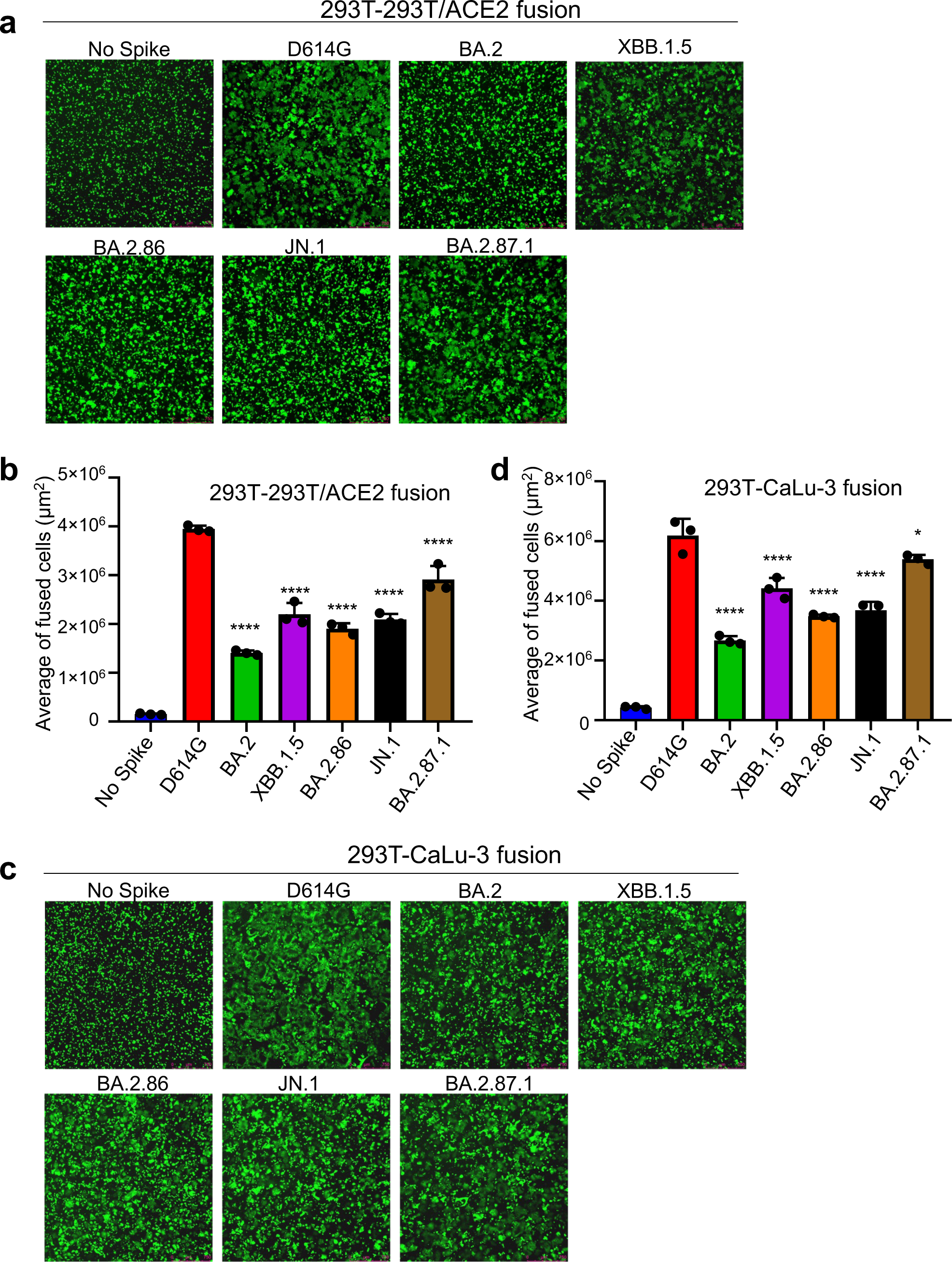
Cell-Cell fusion of BA.2.87.1 and JN.1 spikes alongside other SARS-CoV-2 variants in 293T-ACE2 and CaLu-3 cells. HEK293T cells were co-transfected with plasmids of indicated spikes together with GFP and were cocultured with 293T-ACE2 (a-b) or human lung epithelial CaLu-3 cells (c-d) for 6.5 h (HEK293-ACE2) or 4h (CaLu-3). Cell-cell fusion was imaged and GFP areas of fused cells were quantified (see Methods). D614G and no spike served as positive and negative control, respectively. Comparisons of the extent of cell-cell fusion were made for each Omicron subvariant against D614G. Scale bars represent 150 μM. Bars in (b and d) represent means ± standard error. Dots represent three images from two biological replicates. Statistical significance relative to D614G was determined using a one-way repeated measures ANOVA with Bonferroni’s multiple testing correction (n = 3). p values are displayed as ns p > 0.05, *p < 0.05, and ****p < 0.0001.

We next determined the surface expression level of spike proteins in 293T cells used to produce the lentiviral pseudotyped viruses by flow cytometry. We found that XBB.1.5 exhibited the highest expression, followed by D614G and BA.2.86. Interestingly, BA.2, JN.1, an BA.2.87.1 all exhibited decreased surface expression relative to D614G, with BA.2.87.1 being the lowest (**Figs. 4a-b**). This patten is corroborated by western blotting analysis of the lysate of these producer cells which depicts overall less spike expression for all Omicron variants except for XBB.1.5 (upper panel, **Fig. 4c**). The differences in spike protein expression, including on the plasma membrane, were not due to artifacts of transfection efficiency, given the similar levels of HIV-1 lentiviral Gag expression detected by an anti-P24 antibody (middle panel, **Fig. 4c**) and the comparable signals of GAPDH detected by anti-GAPDH (lower panel, **Fig. 4c)**. Importantly, despite the relatively low level of expression, BA.2.87.1 and JN.1 both exhibited increased processing of spike into the S1 and S2 subunits as compared to the parental D614G and their ancestral BA.2, as quantified by the S1/S and S2/S ratios (**Fig. 4c**).

**FIG 4.**
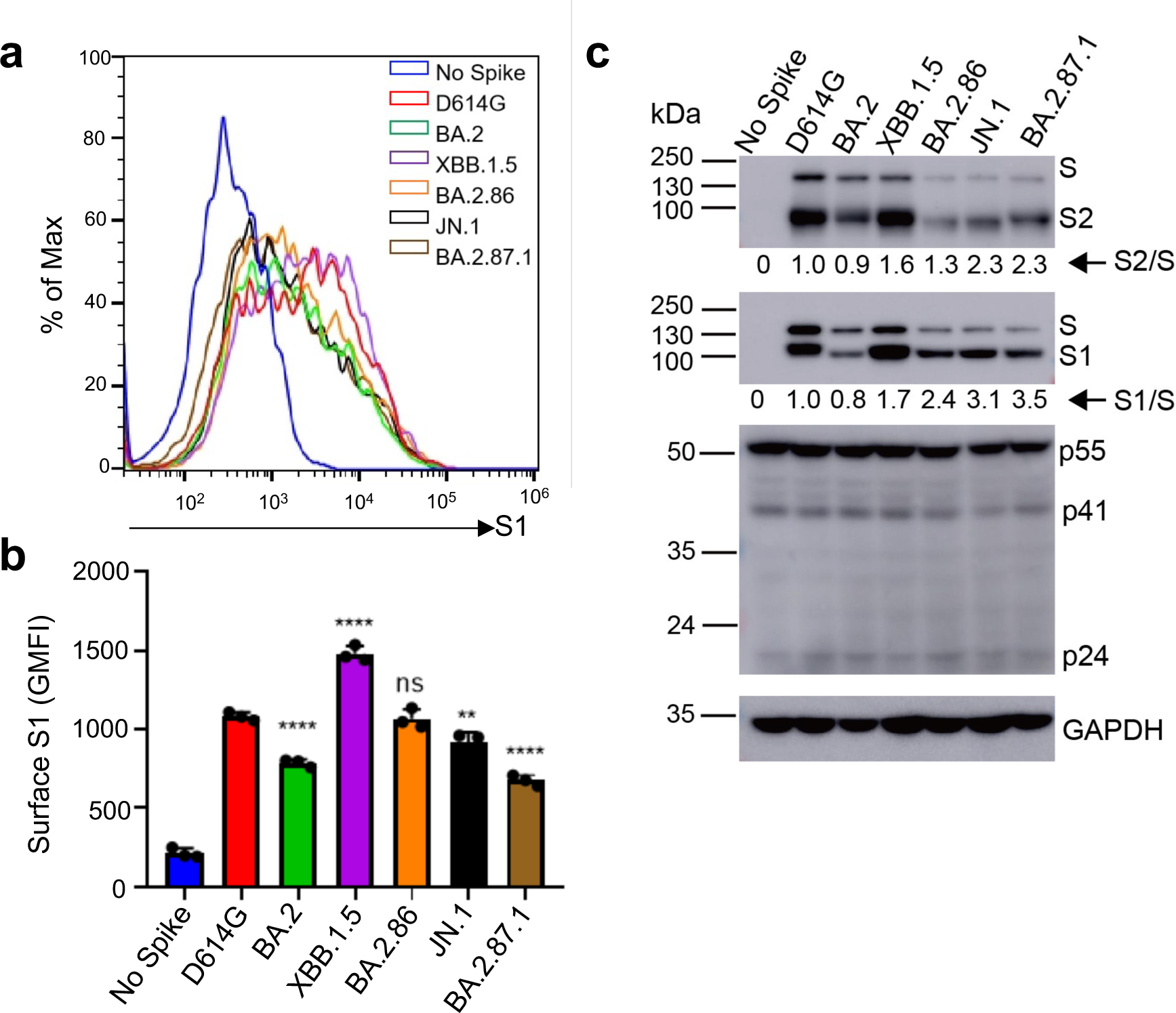
Surface expression and processing of BA.2.87.1, JN.1, and other spike proteins. (a-b) Cell surface expression of spike proteins. HEK293T cells used for production of pseudotyped lentiviral vectors bearing indicated spikes of interest were fixed and stained for spike with an anti-S1 specific antibody T62 followed by flow cytometric analyses. (a) Histogram plots of anti-S1 signals in transfected cells. (b) Mean fluorescence intensities of individual subvariants from (a).(c) Spike expression and processing. HEK293T cells used to produce pseudotyped vectors were lysed and probed with anti-S1, anti-S2, anti-GAPDH, or anti-p24 antibodies; spike processing was quantified using NIH ImageJ to determine the S1/S or S2/S ratio and normalized to D614G (D614G = 1.0). Bars in (b) represent means ± standard error. Dots represent three biological replicates from one typical experiment. Significance relative to D614G was determined using a one-way repeated measures ANOVA with Bonferroni’s multiple testing correction (n = 3). p values are displayed as ns p > 0.05, ** p < 0.01, and ****p < 0.0001.

## DISCUSSION

The continued tracking and characterization of emerging variants of SARS-CoV-2 has proven critical to maintaining pandemic control strategies including vaccination. In accordance with the variants swift rise to dominance, in this work we showed that JN.1 exhibits the lowest nAb titers for both bivalent-vaccinated individuals and first-responders/ICU-admitted COVID-19 patients. The decrease in neutralization titers against JN.1 relative to BA.2.86 is consistent with data published by others, and also explains, at least in part, why JN.1 has become a globally dominant variant compared to its ancestral BA.2.86 (6, 14, 20). Interestingly, we discovered that the newly emerged BA.2.87.1 variant possesses an increased sensitivity to neutralization by these sera compared to JN.1, implying that this variant may not be able to outcompete the current JN.1 and become predominant. However, given that a single L455S mutation in the spike of JN.1 can dramatically increase the nAb evasion of BA.2.86 (3, 14, 21), there is a possibility that additional mutations in BA.2.87.1 could similarly result in new variants that dramatically enhance the nAb escape.

It is currently unclear what amino acid changes in the BA.2.87.1 spike are responsible for the enhanced neutralization by nAb generated by the bivalent mRNA vaccine and JN.1-wave infection. However, given the differences in spike between BA.2.87.1 and others including BA.2 and JN.1 shown in **Fig. 1a**, we speculate that two N-terminal deletions, specifically 15-25del and 136-146del, might have contributed to the comparatively higher nAb titers against BA.2.87.1 compared to BA.2.86/JN.1 and XBB.1.5 — both lacking these deletions. Moreover, BA.2, which serves as the precursor to recent Omicron subvariants and is devoid of these two deletions, demonstrates approximately a 3.2-fold increased titer against BA.2.87.1 (**Fig. 2a**). These findings collectively support for a potential role of these deletions in nAb evasion, which was confirmed by a recent preprint (32). Beyond the N-terminal deletions, the presence of eight amino acid changes in the Receptor Binding Domain (RBD), along with seven amino acid modifications in the furin cleavage site and S2 of the spike (**Fig. 1a**), could alter the ACE2 binding and/or viral membrane fusion capabilities of BA.2.87.1, thus contributing to the varied entry efficiency of BA.2.87.1 (**Fig. 1b-c**). These amino acid changes could also explain the loss of sensitivity of BA.2.87.1 to mAb 2B04 yet re-gain of its neutralizing by S309 (**Fig. 2d** and **Fig. S1e**). Nevertheless, it’s crucial to acknowledge that the replication of BA.2.87.1 diverges from entry mechanisms, and mutations in non-spike regions of the genome could also hold significant roles. Therefore, a comprehensive analysis of the replication of authentic BA.2.87.1 will provide insights into the impact of spike mutations on immune evasion and replication.

In this work, we found that antibodies elicited by BA.2.86/JN.1-wave infection did not effectively neutralize BA.2.86/JN.1 compared to D614G, potentially due to immune imprinting, which has been observed for BA.4/5 and XBB.1.5 variants by ours and other groups (22–24). Immune imprinting arises through two general mechanisms, one is that the immune system prioritizes a recalled response over a new one (“antigenic seniority”), and the other is that new response is actively suppressed (“primary addiction”) (25, 26). Importantly, SARS-CoV-2 infection and vaccination can both cause immune imprinting, resulting in decreased vaccine efficacy (22). For example, vaccinated individuals who had breakthrough infection with different variants mount nAb response primarily towards the wildtype spike protein (9, 10, 14, 18, 21, 26, 27). In this study, all JN.1 patients in the infected cohort had received some doses of vaccine containing the WT spike (**Table S1**). We suspect that this could explain the relatively low titers of these patient sera against JN.1 as compared to D614G (**Fig. 2b** and **Fig. S1b**) (25, 26, 28). A single antigenic exposure to an Omicron subvariant such as JN.1 may not be sufficient to overcome immune imprinting driven by the monovalent WT vaccines (18, 28–32).

The neutralization pattern of XBB.1.5-monovalent-vaccinated hamster sera against BA.2.87.1 is somewhat surprising. These samples exhibited robust titers against XBB.1.5, BA.2.86 and JN.1 yet showed low titers against D614G, which emphasize the need to move away from WT spike-containing vaccines. Interestingly, the titers against BA.2.87.1 were notably lower than those of other Omicron variants in this cohort, raising the possibility that XBB.1.5 monovalent vaccine may not be able to effectively protect against infection by this new variant in SARS-CoV-2 naïve individuals. However, this concern might be diminished, given that a majority of the world population has been vaccinated and/or infected by SARS-CoV-2, unlike the naïve hamsters in this cohort; this hybrid immunity could offer potential broader protection against emerging variants, including JN.1 and BA.2.87.1 (30–32). Indeed, despite JN.1 exhibiting an enhanced ability to evade the COVID-19 vaccine compared to BA.2.86, recent studies (29, 33, 34) have shown that the monovalent XBB.1.5 vaccine can generate effective nAbs against JN.1, contributing to the control of the rapid JN.1 transmission. Unfortunately, we were unable to confirm the result of hamster serum samples in XBB.1.5 monovalent-vaccinated human population with no prior history of exposure to COVID-19 vaccination or SARS-CoV-2 infection, because XBB.1.5 monovalent vaccination is only allowed as booter to those who had been previously vaccinated. In addition, our finding that BA.2.87.1 does not cluster with the other more recent Omicron variants, but instead resembles D614G and BA.2, further highlights the distinctive antigenic nature of BA.2.87.1, underscoring the need to continue monitoring the SARS-CoV-2 variants and updating the COVID-19 vaccines.

In addition to its distinct antigenic phenotype, BA.2.87.1 spike also displayed changes in spike protein biology. Most noticeably, we found that the BA.2.87.1 spike has increased cell-cell fusion and processing as compared to the other Omicron variants including JN.1. While both phenotypes still fall below the levels of D614G, we cannot rule out the possibility that the pathogenicity and/or tissue tropism of this variant may be altered. Experiments using infectious virus to investigate these biological properties will be necessary. Although viral replication fitness is not a focus of this work, it is important to emphasize that differences exist between immunized and immunologically naïve individuals, which can shape the emergence of new SARS-CoV-2 variants and disease pathogenesis. In immunized individuals, viral replication may be controlled more efficiently in immunized subjects due to the quicker and targeted immune response, leading to faster viral clearance and reduced severity of the infection. However, the immune system’s selective pressure in immunized individuals may also drive the evolution of the virus towards variants that can escape immune recognition, although the replication fitness of these escape variants may vary, and they may not always outcompete the original strains in terms of transmissibility or virulence.

## ACKNOWLEDGEMENTS

We thank the members of Liu lab, especially Panke Qu, for helpful discussion. We also thank the Clinical Research Center/Center for Clinical Research Management of The Ohio State University Wexner Medical Center and The Ohio State University College of Medicine in Columbus, Ohio, specifically J. Brandon Massengill, Francesca Madiai, Dina McGowan, Breona Edwards, Evan Long, and Trina Wemlinger, for collection and processing of samples. We thank Tongqing Zhou at NIH for providing the S309 monoclonal antibody. In addition, we thank Sarah Karow, Madison So, Preston So, Daniela Farkas, and Finny Johns in the clinical trials team of The Ohio State University for sample collection and other supports. We thank Moemen Eltobgy for assistance in sample processing. We specially thank Ashish R. Panchal, Soledad Fernandez, Mirela Anghelina, and Patrick Stevens for their assistance in providing the sample information of the first responders and their household contacts. We thank Peng Ru and Lauren Masters for sequencing and Xiaokang Pan for bioinformatic analysis. S.-L.L., D. J., R.J.G., L.J.S. and E.M.O. were supported by the National Cancer Institute of the NIH under award no. U54CA260582. The content is solely the responsibility of the authors and does not necessarily represent the official views of the National Institutes of Health. This work was also supported by a fund provided by an anonymous private donor to OSU. NIH R01AI090060. M.C. was supported by an NIH T32 training grant (T32AI165391). J. L. was supported by NIH R01AI090060. M.C. was supported by an NIH T32 training grant (T32AI165391). J.S.B. was supported by award number grants UL1TR002733 and KL2TR002734 from the National Center for Advancing Translational Sciences. R.J.G. was additionally supported by the Robert J. Anthony Fund for Cardiovascular Research and the JB Cardiovascular Research Fund, and L.J.S. was partially supported by NIH R01 HD095881.

The authors have no competing interests to disclose.

S.-L.L. conceived and directed the project. R.J.G. led the clinical study and experimental design and implementation. P.L. performed all experiments with assistance from Y.L. P.L. performed data processing and analyses with help from J.N.F and Y-.M.Z. C.C.H and J.L. provided the hamster serum samples. D.J. led SARS-CoV-2 variant genotyping and DNA sequencing analyses. C.C., J.S.B, J.C.H., R.M., and R.J.G provided clinical samples and associated information. P.L., J.N.F, and S.-L.L. wrote the paper. J.L., L.J.S., and E.M.O. provided insightful discussion and revision of the manuscript.

---

**Table S1.** Bivalent-vaccinated HCW and BA.2.86/JN.1-wave first responder cohorts.

|  | Bivalent HCWs<br>(n=13) | BA.2.86-JN.1 Wave<br>Patients<br>(n=9) |
| --- | --- | --- |
| **Age in Years at Sample Collection**<br>**[Median (Range)]** | 37 (25-48) | 53(35-78) |
| **Gender** [n (% of Total)] |  |  |
| Male | 8 (61.5%) | 4 (44.4%) |
| Female | 5 (38.5%) | 5 (55.6%) |
| **Sample Collection Window** | Dec. 2022- Jan.2023 | Nov. 2023-Feb.2024 |
| **Vaccine status** [n (% of Total)] |  |  |
| 2-dose Moderna | NA | 3 (11.1%) |
| 1-dose Moderna+1-dose Pfizer bivalent | NA | 1 (11.1%) |
| 3-dose Pfizer +1-dose Moderna bivalent | 1 (7.7%) | 0 |
| 2-dose Pfizer+1-dose Pfizer bivalent | 1 (7.7%) | 0 |
| 4-dose Pfizer+1-dose Pfizer bivalent | 1 (7.7%) | 0 |
| 3-dose Pfizer+1-dose Pfizer bivalent | 3 (23.1%) | 0 |
| 3-dose Moderna+1-dose Moderna bivalent | 6 (46.2%) | 1 (11.1%) |
| 2-dose Moderna+1 Pfizer+1-dose Pfizer bivalent | 1 (7.7%) | 0 |
| 3-dose Moderna+1-dose Pfizer bivalent+1-dose Moderna bivalent | 0 | 1 (11.1%) |
| 2-dose Pfizer +1-dose Moderna | 0 | 1 (11.1%) |
| 1-dose Pfizer | 0 | 1 (11.1%) |
| 3-dose Moderana+1-dose XBB.1.5 Moderna monovalent | 0 | 1 (11.1%) |
| **Sample Collection Timing** [Median (Range)] |  |  |
| Days from last vaccination | NA | 656 (45-898) |
| Days post the bivalent dose for recipients | 66 (23-108) | NA |
| **COVID-19 positive** [n (% of Total)] | 9 (69.2%) | 9 (100%) |
| Days before sample collection [Median (Range)] | 324 (182-994) | 7 (1-10) |
| **Infected Variants** |  |  |
| JN.1/BA.2.86 | 0 | 2 (22.2%) |
| Undetermined | NA | 7 (77.8%) |

## MATERIALS AND METHODS

### Study cohorts

#### Bivalent Vaccinated HCWs (n=13)

These sera were collected from HCWs at the Ohio State Wexner Medical Center that received at least 2 doses of monovalent vaccine (WT) and 1 dose of bivalent vaccine (WT+BA.4/5) under the approved IRB protocols 2020H0228, 2020H0527, and 2017H0292. 11 individuals received 3 doses of monovalent vaccine (Pfizer or Moderna formulations) and 1 bivalent booster dose (Pfizer). 1 person received 4 doses of monovalent vaccine (Pfizer) and 1 bivalent booster dose (Pfizer). 1 person received 2 doses of Pfizer monovalent vaccine and 1 bivalent booster dose (Pfizer). This cohort ranged from 25-48 years of age and included 8 males and 5 females. Blood was collected between 23-108 days post-bivalent booster dose (see details in **Table S1**).

#### ICU patients infected in BA.2.86/JN.1 wave (n=9)

These sera were collected from ICU patients in the OSU Wexner Medical Center or symptomatic participants in the first responder/household contact STOP-COVID cohort who had reverse transcription PCR positivity for SARS-CoV-2 between the dates of 11/23/2024-2/16/2024 during which the BA.2.86/JN.1 variants were predominantly circulating in Columbus, Ohio, U.S (**Table S1**). Samples were collected under the approved IRBs protocols 2020H0527, 2020H0531, 2020H0240, and 2020H0175. Variant type was confirmed in a subset of samples with available nasopharyngeal swabs by SARS-CoV-2 complete genome next-generation sequencing using Artic v5.3.2 (IDT, Coralville, IA) and Artic v4.1 primer sets (Illumina, San Diego, CA).

#### Hamster cohorts vaccinated with monovalent XBB.1.5 vaccine (*n*=15)

Fifteen 4-week-old golden Syrian hamsters (Envigo, Indianapolis, IN) were immunized intranasally with 1.5 x 10^5^ PFU per animal of XBB.1.5 spike-based monovalent vaccine (recombinant mumps virus expressing spike of XBB.1.5). Three weeks later, hamsters were boosted with the same vaccine at the same dose. Blood was collected at week 5 after initial immunization (week 2 after booster immunization).

### Cell lines

The cell lines utilized in this investigation comprised human epithelial kidney cells (HEK293T, ATCC CRL-11268, RRID: CVCL_1926) and HEK293T cells overexpressing human ACE2 (BEI: NR-52511, RRID: CVCL_A7UK). Additionally, we employed the human epithelial lung carcinoma cell line CaLu-3. HEK293T cell lines were cultured in DMEM Gibco (11965–092) supplemented with 10% fetal bovine serum (Sigma, F1051) and 0.5% penicillin/streptomycin (HyClone, SV30010). CaLu-3 cells (RRID: CVCL_0609) were cultured in EMEM (ATCC, 30-2003) under the same conditions. Cell cultures were maintained at 37°C with 5.0% CO2 and sub-cultured by washing with PBS (Sigma, D5652-10X1L) followed by detachment using 0.05% trypsin + 0.53 mM EDTA (Corning, 25-052-CI).

### Plasmids

All spike constructs are encoded within the pcDNA3.1 backbone and flanked by C-terminal FLAG tags. They were cloned using KpnI and EcoRI restriction sites. D614G, BA.2, BA.2.86, and BA.2.87.1 plasmids were all synthesized by GenScript Biotech (Piscataway, NJ). The BA.2.87.1 spike sequence was generated based on the consensus of the first few reported Isolates: hCoV-19/SouthAfrica/NICD-R13200/2023 EPI_ISL_18849984; hCoV 19/SouthAfrica/NICD-N56614/2023 EPI_ISL_18849985; hCoV-19/SouthAfrica/NICD-N56836/2023 EPI_ISL_18849986; hCoV-19/SouthAfrica/NICD-N57176/2023 EPI_ISL_18849987; hCoV-19/SouthAfrica/NICD-N57208/2023 EPI_ISL_18849988; hCoV-19/SouthAfrica/NICD-N57216/2023 EPI_ISL_18849989; hCoV-19/SouthAfrica/NICD-N57440/2023 EPI_ISL_18849990; hCoV-19/SouthAfrica/NICD-N57469/2023 EPI_ISL_18849991;hCoV-19/South Africa/NICD-R13515/2023 EPI_ISL_18845398; while XBB.1.5 and JN.1 were generated through site-directed mutagenesis of XBB and BA.2.86, respectively. The lentiviral vector used was an HIV-1 based vector called Pnl4-3 with an Env deletion that encodes a *Gaussia luciferase* reporter gene.

### Pseudotyped lentiviral production and infectivity

Pseudotyped lentiviral vectors were generated following established protocols. Briefly, 293T cells were co-transfected using PEI (Transporter 5 Transfection Reagent, Polysciences) at a 2:1 ratio with the Pnl4-3_inGluc vector and the spike plasmid under investigation. Pseudovirus was harvested by collecting media from the cells at 48 and 72 hours post-transfection. The media was then clarified by centrifugation, and equal volumes were utilized to infect the target cells. Luciferase activity was measured by combining 20Ul of infected cell culture media with 20Ul of Gaussia luciferase substrate (0.1 M Tris Ph 7.4, 0.3 M sodium ascorbate, 10 µM coelenterazine) and immediately quantifying luminescence using a BioTek Cytation plate reader. These values were normalized relative to D614G, with D614G set as 1.

### Virus neutralization assay

The pseudotyped lentiviral vector neutralization assay was performed as described previously (10). Briefly, sera samples are serially diluted 4-fold at a starting dilution of 1:40 for 5 total dilutions (1:40, 1:160, 1:640, 1:2560, 1:10240), with one well left without sera. Pseudotyped viruses are diluted based on infectivity readouts in order to normalize them then placed in equal volumes on the diluted sera and incubated 1 hour at 37°C. The sera/virus mixture is then used to infect 293T-ACE2 cells. As described for infectivity, luminescence readouts are collected at 48 and 72 hours and used to determine a neutralization titer at 50% (NT_50_) using least squares fit non-linear regression normalized to the no serum value using GraphPad Prism 9 (San Diego, CA).

### Cell-cell fusion

Direct spike-mediated cell to cell fusion assays were performed by first co-transfecting 293T cells with spike and GFP. 293T cells were incubated 24 hours then detached and reseeded in a plate containing one of two detached target cells; 293T-ACE2 or CaLu-3. 293T-ACE2 cells were incubated for 6.5 hours and CaLu-3 cells 4 hours then fusion was imaged using a Leica Dmi8 microscope. Areas of fusion were quantified using the Leica X Applications Suite software to outline the edges of fields of GFP and quantify then areas. Three images from duplicate wells were randomly taken. Scale bars represent 150 µM and one representative image was selected for presentation.

### Syncytia formation assay

To validate the cell-cell fusion results, a syncytia formation assay was also performed. 293T-ACE2 cells were co-transfected with the spike of interest and GFP and incubated 24 hours before imaging syncytia using a Leica Dmi8 microscope. The images were processed and displayed the same way as the cell-cell fusion results.

### Spike protein surface expression detected by flow cytometry

A portion of 293T cells used to produce the lentiviral vectors were collected by detaching with PBS + 5Mm EDTA and fixed in 3.7% formaldehyde for 10 minutes and room temperature. Cells were then stained with polyclonal anti-SARS-CoV-2 S1 antibody (Sino Biological, 40591-T62; RRID: AB_2893171) followed by anti-Rabbit-IgG-FITC (Sigma, F9887, RRID: AB_259816) secondary to visualize on a Life Technologies Attune NxT flow cytometer. FlowJo v10 (Ashland, OR) is used to analyze data.

### Spike protein processing

The remaining 293T cells used to produce lentiviral vectors are lysed in RIPA buffer (Sigma-Aldrich, R0278) supplemented with protease inhibitor (Sigma, P8340) for 40 minutes on ice. Lysate is collected and a portion is used for SDS-PAGE on a 10% poly-acrylamide gel and transferred to a PVDF membrane for western blotting. Blots were probed with polyclonal anti-SARS-CoV-2 S1 (Sino Biological, 40591-T62; RRID:AB_2893171), anti-S2 (Sino Biological, 40590; RRID:AB_2857932), anti-p24 (NIH HIV Reagent Program, ARP-1513), and anti-GAPDH (Santa Cruz, Cat# sc-47724, RRID: AB_627678). Secondary antibodies used were Anti-Rabbit-IgG-HRP (Sigma, A9169; RRID:AB_258434) and Anti-Mouse (Sigma, Cat# A5278, RRID: AB_258232). Blots were visualized via Immobilon Crescendo Western HRP substrate (Millipore, WBLUR0500) and exposed on a GE Amersham Imager 600. Band intensities were quantified using NIH Image J analysis software (Bethesda, MD).

### Antigenic mapping

Antigenic cartography was performed using the Racmacs program (v1.1.35) by following instructions provided on their GitHub (https://github.com/acorg/Racmacs/tree/master). Briefly, the program is run in R (Vienna, Austria) and works by taking raw neutralization titers and log2 transforming them to create a distance table for the individual antigens (spike protein) and sera samples. The program then uses this table to perform multidimensional scaling to plot the individual antigen and sera samples as single points where distance between the points directly correlates to antigenic differences. 1 antigenic distance unit (AU), represented by one side of a square in the plots, is equivalent to a 2-fold change in neutralization titers. Optimization settings were kept on default (2 dimensions, 500 optimizations, minimum column basis “none”). Maps were saved as images via the “view(map)” function and labeled using Microsoft Office PowerPoint.

### Statistical analysis

Statistical analyses were conducted using GraphPad Prism 9. Error bars in the figures represent means with standard error. In Figs 1b and 1c, Figs. 3b and 3d, Fig. 4b, and Fig. S2b, comparisons between viruses were made using a one-way ANOVA with Bonferroni post-test. Neutralization titers were determined using least-squares non-linear regression. In Figs 2a-c, error bars represent geometric means with 95% confidence intervals. Comparisons between viruses in these figures were made using a repeated measures one-way ANOVA with Bonferroni post-test. To better approximate normality, comparisons were conducted using log10 transformed NT_50_ values. Error bars in Fig. 2d represent means ± standard deviation. Cell-cell fusion and syncytia formation shown in Figs. 3a, 3c, and Fig. S2a was quantified using the Leica X Applications Suite software. Spike processing shown in Fig. 4c was quantified by NIH ImageJ; the values are then set relative to D614G, with D614G = 1.0.

## RESOURCE AVAILABILITY

Data reported in this paper will be shared by the lead contact upon request, Dr. Shan-Lu Liu. Any additional information required to reanalyze the data reported in this paper is available from the lead contact upon request.

## REFERENCES

1. Rambaut A, Holmes EC, O’Toole Á, Hill V, McCrone JT, Ruis C, du Plessis L, Pybus OG. 2020. A dynamic nomenclature proposal for SARS-CoV-2 lineages to assist genomic epidemiology. Nature Microbiology 5:1403–1407.

2. CDC. 2022. COVID Data Tracker, Variant Proportions.

3. Wang Q, Guo Y, Liu L, Schwanz LT, Li Z, Nair MS, Ho J, Zhang RM, Iketani S, Yu J, Huang Y, Qu Y, Valdez R, Lauring AS, Huang Y, Gordon A, Wang HH, Liu L, Ho DD. 2023. Antigenicity and receptor affinity of SARS-CoV-2 BA.2.86 spike. Nature 624:639–644.

4. Qu P, Xu K, Faraone JN, Goodarzi N, Zheng Y-M, Carlin C, Bednash JS, Horowitz JC, Mallampalli RK, Saif LJ, Oltz EM, Jones D, Gumina RJ, Liu S-L. 2024. Immune evasion, infectivity, and fusogenicity of SARS-CoV-2 BA.2.86 and FLip variants. Cell 187:585–595.e6.

5. Liu Z, Zhou J, Wang W, Zhang G, Xing L, Zhang K, Wang Y, Xu W, Wang Q, Man Q, Wang Q, Ying T, Zhu Y, Jiang S, Lu L. 2024. Neutralization of SARS-CoV-2 BA.2.86 and JN.1 by CF501 adjuvant-enhanced immune responses targeting the conserved epitopes in ancestral RBD. Cell Rep Med doi:10.1016/j.xcrm.2024.101445:101445.

6. Yang S, Yu Y, Xu Y, Jian F, Song W, Yisimayi A, Wang P, Wang J, Liu J, Yu L, Niu X, Wang J, Wang Y, Shao F, Jin R, Wang Y, Cao Y. 2024. Fast evolution of SARS-CoV-2 BA.2.86 to JN.1 under heavy immune pressure. Lancet Infect Dis 24:e70–e72.

7. CDC. 2/9/2024 2024. CDC Tracks New SARS-CoV-2 Variant, BA.2.87.1.

8. Zeng C, Evans JP, Qu P, Faraone J, Zheng YM, Carlin C, Bednash JS, Zhou T, Lozanski G, Mallampalli R, Saif LJ, Oltz EM, Mohler P, Xu K, Gumina RJ, Liu SL. 2021. Neutralization and Stability of SARS-CoV-2 Omicron Variant. bioRxiv doi:10.1101/2021.12.16.472934.

9. Qu P, Faraone JN, Evans JP, Zheng Y-M, Carlin C, Anghelina M, Stevens P, Fernandez S, Jones D, Panchal AR, Saif LJ, Oltz EM, Zhang B, Zhou T, Xu K, Gumina RJ, Liu S-L. 2023. Enhanced evasion of neutralizing antibody response by Omicron XBB.1.5, CH.1.1, and CA.3.1 variants. Cell Reports 42:112443.

10. Faraone JN, Qu P, Goodarzi N, Zheng Y-M, Carlin C, Saif LJ, Oltz EM, Xu K, Jones D, Gumina RJ, Liu S-L. 2023. Immune Evasion and Membrane Fusion of SARS-CoV-2 XBB Subvariants EG.5.1 and XBB.2.3. Emerging Microbes and Infections 12(2):2270069.

11. Evans JP, Zeng C, Qu P, Faraone J, Zheng YM, Carlin C, Bednash JS, Zhou T, Lozanski G, Mallampalli R, Saif LJ, Oltz EM, Mohler PJ, Xu K, Gumina RJ, Liu SL. 2022. Neutralization of SARS-CoV-2 Omicron sub-lineages BA.1, BA.1.1, and BA.2. Cell Host Microbe 30:1093–1102.e3.

12. Qu P, Evans JP, Faraone JN, Zheng YM, Carlin C, Anghelina M, Stevens P, Fernandez S, Jones D, Lozanski G, Panchal A, Saif LJ, Oltz EM, Xu K, Gumina RJ, Liu SL. 2023. Enhanced neutralization resistance of SARS-CoV-2 Omicron subvariants BQ.1, BQ.1.1, BA.4.6, BF.7, and BA.2.75.2. Cell Host Microbe 31:9–17.e3.

13. Zeng C, Evans JP, Pearson R, Qu P, Zheng YM, Robinson RT, Hall-Stoodley L, Yount J, Pannu S, Mallampalli RK, Saif L, Oltz E, Lozanski G, Liu SL. 2020. Neutralizing antibody against SARS-CoV-2 spike in COVID-19 patients, health care workers, and convalescent plasma donors. JCI Insight 5.

14. Zhang L, Kempf A, Nehlmeier I, Cossmann A, Richter A, Bdeir N, Graichen L, Moldenhauer AS, Dopfer-Jablonka A, Stankov MV, Simon-Loriere E, Schulz SR, Jäck HM, Čičin-Šain L, Behrens GMN, Drosten C, Hoffmann M, Pöhlmann S. 2024. SARS-CoV-2 BA.2.86 enters lung cells and evades neutralizing antibodies with high efficiency. Cell 187:596–608.e17.

15. Wang X, Lu L, Jiang S. 2024. SARS-CoV-2 evolution from the BA.2.86 to JN.1 variants: unexpected consequences. Trends Immunol 45:81–84.

16. Chen RE, Winkler ES, Case JB, Aziati ID, Bricker TL, Joshi A, Darling TL, Ying B, Errico JM, Shrihari S, VanBlargan LA, Xie X, Gilchuk P, Zost SJ, Droit L, Liu Z, Stumpf S, Wang D, Handley SA, Stine WB, Jr., Shi PY, Davis-Gardner ME, Suthar MS, Knight MG, Andino R, Chiu CY, Ellebedy AH, Fremont DH, Whelan SPJ, Crowe JE, Jr., Purcell L, Corti D, Boon ACM, Diamond MS. 2021. In vivo monoclonal antibody efficacy against SARS-CoV-2 variant strains. Nature 596:103–108.

17. Errico JM, Zhao H, Chen RE, Liu Z, Case JB, Ma M, Schmitz AJ, Rau MJ, Fitzpatrick JAJ, Shi PY, Diamond MS, Whelan SPJ, Ellebedy AH, Fremont DH. 2021. Structural mechanism of SARS-CoV-2 neutralization by two murine antibodies targeting the RBD. Cell Reps 37:109881.

18. Yang S, Yu Y, Jian F, Song W, Yisimayi A, Chen X, Xu Y, Wang P, Wang J, Yu L, Niu X, Wang J, Xiao T, An R, Wang Y, Gu Q, Shao F, Jin R, Shen Z, Wang Y, Cao Y. 2023. Antigenicity and infectivity characterisation of SARS-CoV-2 BA.2.86. Lancet Infect Dis 23:e457–e459.

19. Alsoussi WB, Turner JS, Case JB, Zhao H, Schmitz AJ, Zhou JQ, Chen RE, Lei T, Rizk AA, McIntire KM, Winkler ES, Fox JM, Kafai NM, Thackray LB, Hassan AO, Amanat F, Krammer F, Watson CT, Kleinstein SH, Fremont DH, Diamond MS, Ellebedy AH. 2020. A Potently Neutralizing Antibody Protects Mice against SARS-CoV-2 Infection. J Immunol 205:915–922.

20. Uriu K, Ito J, Kosugi Y, Tanaka YL, Mugita Y, Guo Z, Hinay AA, Jr., Putri O, Kim Y, Shimizu R, Begum MM, Jonathan M, Saito A, Ikeda T, Sato K. 2023. Transmissibility, infectivity, and immune evasion of the SARS-CoV-2 BA.2.86 variant. Lancet Infect Dis 23:e460–e461.

21. Tamura T, Mizuma K, Nasser H, Deguchi S, Padilla-Blanco M, Oda Y, Uriu K, Tolentino JEM, Tsujino S, Suzuki R, Kojima I, Nao N, Shimizu R, Wang L, Tsuda M, Jonathan M, Kosugi Y, Guo Z, Hinay AA, Jr., Putri O, Kim Y, Tanaka YL, Asakura H, Nagashima M, Sadamasu K, Yoshimura K, Saito A, Ito J, Irie T, Tanaka S, Zahradnik J, Ikeda T, Takayama K, Matsuno K, Fukuhara T, Sato K. 2024. Virological characteristics of the SARS-CoV-2 BA.2.86 variant. Cell Host Microbe 32:170–180.e12.

22. Addetia A, Piccoli L, Case JB, Park YJ, Beltramello M, Guarino B, Dang H, de Melo GD, Pinto D, Sprouse K, Scheaffer SM, Bassi J, Silacci-Fregni C, Muoio F, Dini M, Vincenzetti L, Acosta R, Johnson D, Subramanian S, Saliba C, Giurdanella M, Lombardo G, Leoni G, Culap K, McAlister C, Rajesh A, Dellota E, Jr., Zhou J, Farhat N, Bohan D, Noack J, Chen A, Lempp FA, Quispe J, Kergoat L, Larrous F, Cameroni E, Whitener B, Giannini O, Cippà P, Ceschi A, Ferrari P, Franzetti-Pellanda A, Biggiogero M, Garzoni C, Zappi S, Bernasconi L, Kim MJ, Rosen LE, Schnell G, et al. 2023. Neutralization, effector function and immune imprinting of Omicron variants. Nature 621:592–601.

23. Reynolds CJ, Pade C, Gibbons JM, Otter AD, Lin KM, Muñoz Sandoval D, Pieper FP, Butler DK, Liu S, Joy G, Forooghi N, Treibel TA, Manisty C, Moon JC, Semper A, Brooks T, McKnight Á, Altmann DM, Boyton RJ, Abbass H, Abiodun A, Alfarih M, Alldis Z, Altmann DM, Amin OE, Andiapen M, Artico J, Augusto JB, Baca GL, Bailey SNL, Bhuva AN, Boulter A, Bowles R, Boyton RJ, Bracken OV, O’Brien B, Brooks T, Bullock N, Butler DK, Captur G, Carr O, Champion N, Chan C, Chandran A, Coleman T, Couto de Sousa J, Couto-Parada X, Cross E, Cutino-Moguel T, D’Arcangelo S, et al. 2022. Immune boosting by B.1.1.529 (Omicron) depends on previous SARS-CoV-2 exposure. Science 377:eabq1841.

24. Yisimayi A, Song W, Wang J, Jian F, Yu Y, Chen X, Xu Y, Yang S, Niu X, Xiao T, Wang J, Zhao L, Sun H, An R, Zhang N, Wang Y, Wang P, Yu L, Lv Z, Gu Q, Shao F, Jin R, Shen Z, Xie XS, Wang Y, Cao Y. 2024. Repeated Omicron exposures override ancestral SARS-CoV-2 immune imprinting. Nature 625:148–156.

25. Faraone JN, Liu SL. 2023. Immune imprinting as a barrier to effective COVID-19 vaccines. Cell Rep Med 4:101291.

26. Wang Q, Guo Y, Tam AR, Valdez R, Gordon A, Liu L, Ho DD. 2023. Deep immunological imprinting due to the ancestral spike in the current bivalent COVID-19 vaccine. Cell Reports Medicine.

27. Qu P, Faraone J, Evans JP, Zou X, Zheng YM, Carlin C, Bednash JS, Lozanski G, Mallampalli RK, Saif LJ, Oltz EM, Mohler PJ, Gumina RJ, Liu SL. 2022. Neutralization of the SARS-CoV-2 Omicron BA.4/5 and BA.2.12.1 Subvariants. N Engl J Med 386:2526–2528.

28. Yisimayi A, Song W, Wang J, Jian F, Yu Y, Chen X, Xu Y, Yang S, Niu X, Xiao T, Wang J, Zhao L, Sun H, An R, Zhang N, Wang Y, Wang P, Yu L, Gu Q, Shao F, Jin R, Shen Z, Xie XS, Wang Y, Cao Y. 2023. Repeated Omicron infection alleviates SARS-CoV-2 immune imprinting. bioRxiv doi:10.1101/2023.05.01.538516:2023.05.01.538516.

29. Wang Q, Guo Y, Bowen A, Mellis IA, Valdez R, Gherasim C, Gordon A, Liu L, Ho DD. 2024. XBB.1.5 monovalent mRNA vaccine booster elicits robust neutralizing antibodies against XBB subvariants and JN.1. Cell Host Microbe doi:10.1016/j.chom.2024.01.014.

30. Wang X, Jiang S, Ma W, Li C, Liu C, Xie F, Zhu J, Zhan Y, Jiang S, Li M, Zhang Y, Wang P. 2024. Robust Neutralization of SARS-CoV-2 Variants Including JN.1 and BA.2.87.1 by Trivalent XBB Vaccine-Induced Antibodies. bioRxiv doi:10.1101/2024.02.16.580615:2024.02.16.580615.

31. Lasrado N, Rössler A, Rowe M, Collier A-rY, Barouch DH. 2024. Neutralization of SARS-CoV-2 Omicron subvariant BA.2.87.1. Vaccine 10.1016/j.vaccine.2024.03.007.

32. Yang S, Yu Y, Jian F, Yisimayi A, Song W, Liu J, Wang P, Xu Y, Wang J, Niu X, Yu L, Wang Y, Shao F, Jin R, Wang Y, Cao Y. 2024. Antigenicity assessment of SARS-CoV-2 saltation variant BA.2.87.1. bioRxiv doi:10.1101/2024.03.07.583823 %J bioRxiv:2024.03.07.583823.

33. Chalkias S, McGhee N, Whatley JL, Essink B, Brosz A, Tomassini JE, Girard B, Edwards DK, Wu K, Nasir A, Lee D, Avena LE, Feng J, Deng W, Montefiori DC, Baden LR, Miller JM, Das R. 2024. Interim report of the reactogenicity and immunogenicity of SARS-CoV-2 XBB-containing vaccines. J Infect Dis doi:10.1093/infdis/jiae067.

34. Jain S, Kumar S, Lai L, Linderman S, Malik AA, Ellis ML, Godbole S, Solis D, Sahoo MK, Bechnak K, Paredes I, Tanios R, Kazzi B, Dib SM, Litvack MB, Wimalasena ST, Ciric C, Rostad C, West R, Teng IT, Wang D, Edupuganti S, Kwong PD, Rouphael N, Pinsky BA, Douek DC, Wrammert J, Moreno A, Suthar MS. 2024. XBB.1.5 monovalent booster improves antibody binding and neutralization against emerging SARS-CoV-2 Omicron variants. bioRxiv doi:10.1101/2024.02.03.578771.

