## Supplementary Figures 1 and 2 for "Distinct Patterns of SARS-CoV-2 BA.2.87.1 and JN.1 Variants in Immune Evasion, Antigenicity and Cell-Cell Fusion"

a

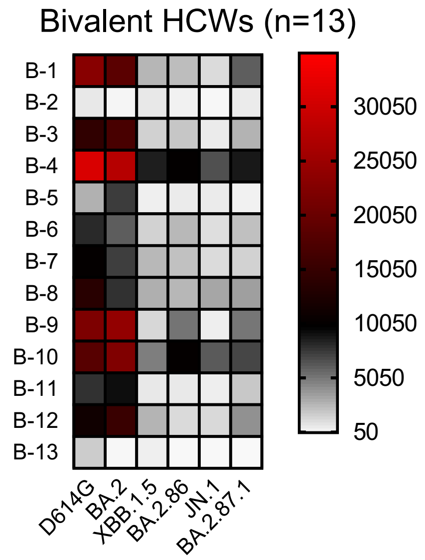


b

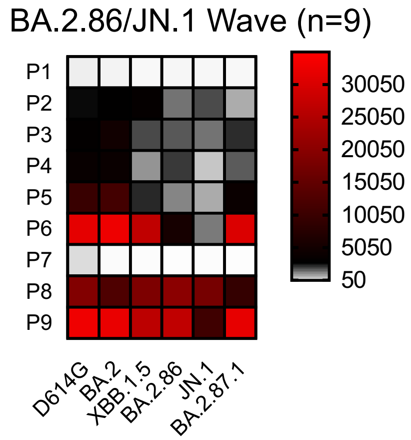


c

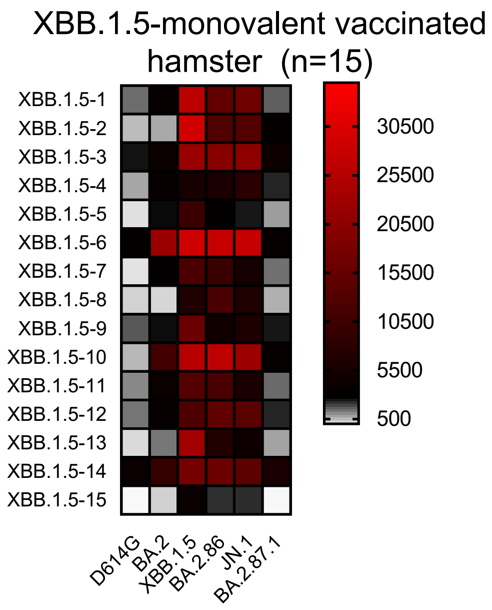


d

| Variants | **S309** IC50($\mu$g/ml) | **2B04** IC50($\mu$g/ml) |
|---|---|---|
| D614G | 0.13±0.03 | 0.01±0.00 |
| BA.2 | 0.41±0.08 | >12 |
| XBB.1.5 | 0.30±0.13 | >12 |
| BA.2.86 | 6.22±1.35 | >12 |
| JN.1 | >12 | >12 |
| BA.2.87.1 | 0.62±0.30 | >12 |

e

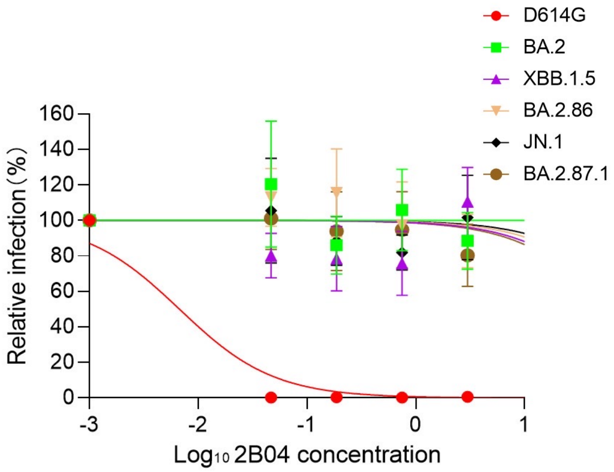


**FIG S1 Heatmap display of nAb escape in three cohorts.** NAb tiers against BA.2.87.1, BA.2.86, JN.1, XBB.1.5, BA.2 and ancestral D614G shown in Figs. 2a-c are respectively displayed as heat maps for sera from health care workers (HCWs) (n = 13) (a), for sera from Columbus first-responder/household contact cohort (P1 to P5) and ICU patients admitted to OSU Wexner Medical Center (P6 to P9) (total n=9) during when the BA.2.86/JN.1 variants were predominantly circulating in Columbus, Ohio (b), and for sera from Golden Syrian hamsters inoculated with 2 doses of monovalent XBB.1.5 vaccine (1.5 x $10^5$ PFU per hamster, n=15), with blood being collected 5 weeks after inoculation (c). The neutralization assays were also performed to assess the effectiveness of mAbs S309 and 2B04 in neutralizing SARS-CoV-2 variants (d-e). The calculated IC50 values (means ± standard deviation are shown (d). The neutralization curves of 2B04 were shown in (e).

FIG S1

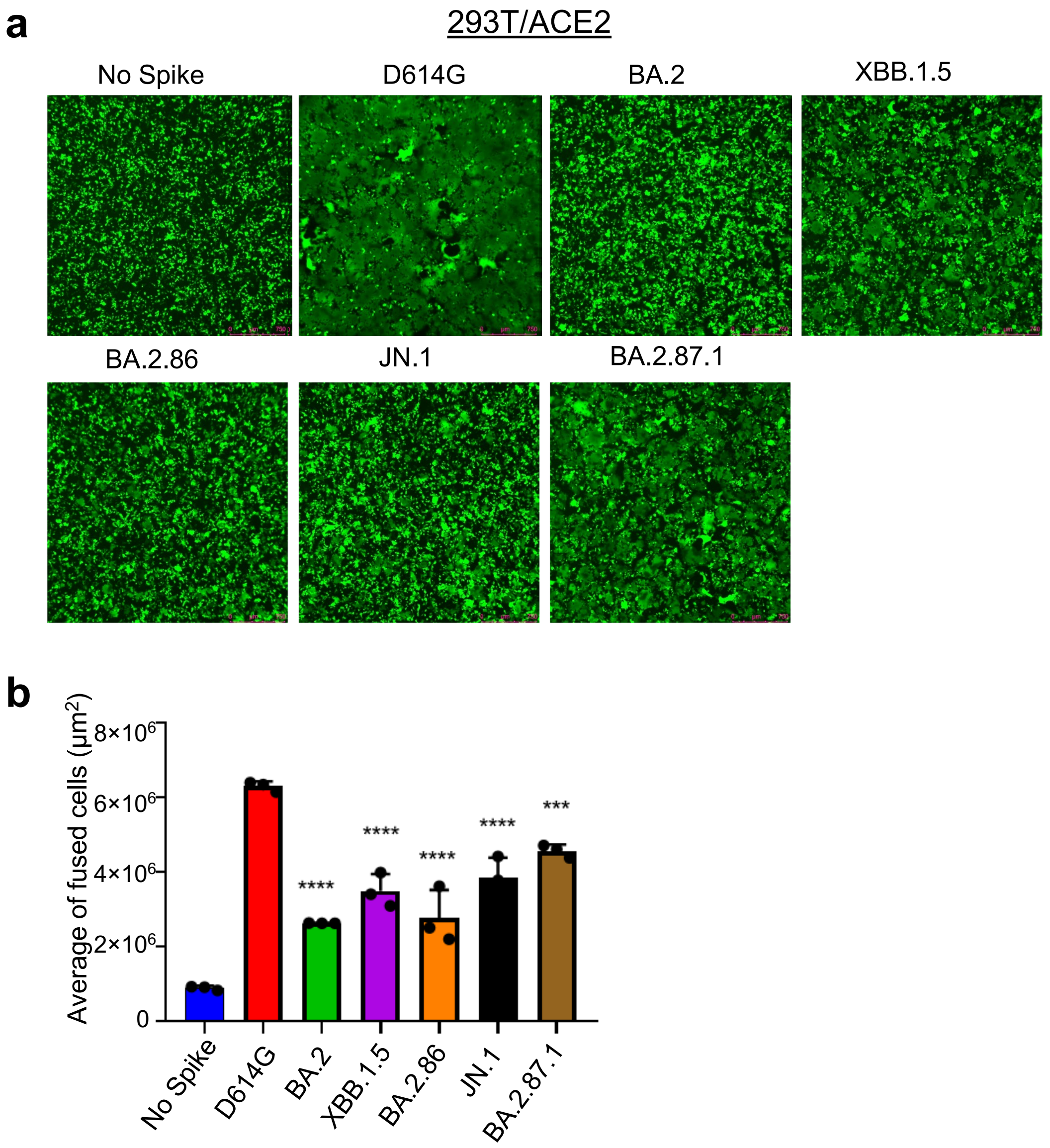


**FIG S2 Syncytia formation induced by BA.2.87.1, JN.1 or other Omicron spike proteins.** (a) 293T-ACE2 cells were transfected to produce the spikes of interest along with GFP and incubated 24 hours before imaging fusion. (b) GFP areas of fused cells were quantified (see Methods). D614G and no spike were included as positive and negative control, respectively. Comparisons in extents of syncytia formation for each Omicron subvariant were made against D614G, with scale bars representing 150 μM. Bars in (b) represent means ± standard error. Statistical significance relative to D614G was determined using a one-way repeated measures ANOVA with Bonferroni's multiple testing correction ($n = 3$). p values are displayed as ***$p < 0.001$, and ****$p < 0.0001$.
